# Cell-type and endocannabinoid specific synapse connectivity in the adult nucleus accumbens core

**DOI:** 10.1101/613497

**Authors:** Marion A. Deroche, Olivier Lassalle, Olivier J. Manzoni

## Abstract

The nucleus accumbens (NAc) is a mesocorticolimbic structure that integrates cognitive, emotional and motor functions. Although its role in psychiatric disorders is widely acknowledged, the understanding of its circuitry is not complete. Here we combined optogenetic and whole-cell recordings to draw a functional portrait of excitatory disambiguated synapses onto D1 and D2 medium spiny neurons (MSNs) in the adult mouse NAc core. Comparing synaptic properties of ventral hippocampus (vHipp), basolateral amygdala (BLA) and prefrontal cortex (PFC) inputs revealed a hierarchy of synaptic inputs and feedforward inhibition that depends on the identity of the postsynaptic target MSN. Thus, the BLA is the dominant excitatory pathway onto D1 MSNs (BLA > PFC = vHipp) while PFC inputs dominate D2 MSNs (PFC > vHipp > BLA). Feedforward inhibition of MSN firing too, was input and cell-type specific: while minimal at vHipp-D1 and vHipp-D2 inputs; it inhibited with similar efficacy BLA-D1 or BLA-D2 inputs, was minimal at PFC-D1 but maximal at PFC-D2 inputs. We also tested the hypothesis that endocannabinoids endow excitatory circuits with pathway- and cell-specific plasticity. Thus, while CB1 receptors (CB1R) uniformly depress excitatory pathways irrespective of MSNs identity, TRPV1 receptors (TRPV1R) bidirectionally control inputs onto the NAc core in a pathway-specific manner. Finally, we show how the interplay of TRPV1R/CB1R shapes plasticity at identified BLA-NAc synapses. Together these data shed new light on synapse and circuit specificity in the adult NAc core and illustrate how endocannabinoids contribute to pathway-specific synaptic plasticity.

**SIGNIFICANCE STATEMENT:** We examined the impact of connections from the ventral hippocampus (vHipp,) basolateral amygdala (BLA) and prefrontal cortex (PFC) onto identified medium spiny neurons (MSN) in the adult accumbens core. We found BLA inputs were strongest at D1 MSNs while PFC inputs dominate D2 MSNs. We evaluated the role of the endocannabinoid system in pathway- and cell-specific plasticity and found that CB1 receptors (CB1R) and TRPV1 receptors (TRPV1R) bidirectionally control synaptic transmission and plasticity onto accumbens’ MSNs in a pathway- and cell-specific manner. Finally, we clarify how the interplay of TRPV1R/CB1R shapes plasticity at identified BLA-NAc synapses.

## INTRODUCTION

The nucleus accumbens (NAc) is a mesocorticolimbic structure (Humphries and Prescott, 2010) that integrates cognitive, emotional and motor functions (Floresco, 2015). Although the NAc’s role in neurological and psychiatric disorders including anxiety, depression, addiction and intellectual disability (Goto and Grace, 2008; Kasanetz et al., 2010; Sesack and Grace, 2010; Lafourcade et al., 2011; Jung et al., 2012; Neuhofer et al., 2015, 2018; Bosch-Bouju et al., 2016; Manduca et al., 2017) is widely acknowledged (Salgado and Kaplitt, 2015), a detailed understanding of its physiological mechanisms is lacking.

The NAc consists of at least two subregions, a medial “shell” region and a more lateral “core” component (Zahm and Brog, 1992). The principal cell type is GABAergic projection medium-spiny neurons (MSNs) which express either D1 or D2 receptors and play specific roles in NAc-mediated behaviors and disorders (Lobo and Nestler, 2011; Francis et al., 2015).

In young mice, MSNs’ intrinsic and synaptic properties diverge: D2 MSNs are more excitable than D1. Whether these specific differences in MSNs persist in adulthood remains unknown. NAc MSNs receive and integrate glutamatergic inputs, most notably from the prelimbic region of the prefrontal cortex (PFC), the ventral hippocampus (vHipp) and basolateral amygdala (BLA) (Groenewegen et al., 1999; Britt et al., 2012) but also from the thalamus, dorsal hippocampus, VTA and insular cortex (Stratford and Wirtshafter, 2013; Qi et al., 2016; Zhu et al., 2016; Trouche et al., 2018; Rogers-Carter et al., 2019). These regions process dissociable types of information and the specific activation of these pathways can elicit distinct behavioral functions via interactions with the NAc (Goto and Grace, 2008; Sesack and Grace, 2010). The PFC and the NAc interact in behaviors that require executive attention or working memory (Christakou et al., 2001, 2004; Cools et al., 2007), that place high demands on attention (Christakou et al., 2004) or necessitate to link behaviors across contexts (Floresco et al., 1999). The vHipp-NAc pathway is essential for spatial navigation in relation to goal direct behavior (Floresco et al., 1997; Ito et al., 2008; Mannella et al., 2013) and in encoding the temporal dynamics of decision making (Cardinal and Howes, 2005; Eichenbaum, 2014; Abela et al., 2015). By contrast, the BLA-NAc pathway plays a large role in conditioned emotional responses (Everitt et al., 2003; LeDoux, 2003; Tye et al., 2011; Beyeler et al., 2016, 2018) and in forming associations between stimuli that predict appetitive or aversive consequences (Everitt et al., 1991; McLaughlin and Floresco, 2007; Shiflett and Balleine, 2010; Fernando et al., 2013).

The mesocorticolimbic endocannabinoid (eCB) system modulates a vast array of synaptic functions (Robbe et al., 2002; Lafourcade et al., 2007; Araque et al., 2017). eCB-mediated long-term depression was originally discovered in the NAc/ventral striatum (Robbe et al., 2002) and dorsal striatum (Gerdeman et al., 2002). eCB dysfunction is implicated as a major causal factor in a plethora of synaptopathies linked to the NAc (Kasanetz et al., 2010; Lafourcade et al., 2011; Jung et al., 2012; Neuhofer et al., 2015, 2018; Bosch-Bouju et al., 2016; Araque et al., 2017; Manduca et al., 2017). In the dorsal striatum, chemically induced eCB-synaptic plasticity shows pathway but not MSN subtype specificity (Wu et al., 2015). Whether eCBs participate to pathway-specificity in the NAc and a fortiori the mechanistic underpinnings, is unknown. Here, we explored the innervation and synaptic properties of PFC, BLA and vHipp to the NAc core. In addition, we assayed each pathway for eCB receptors and dissected eCB-plasticity at BLA inputs. We report that adult D1- are inherently more excitable than D2 MSNs and that the hierarchy of excitatory inputs depends on the identity of the postsynaptic target MSN and on circuit specific feedforward inhibition. Finally, we provide evidence that the eCB system endows excitatory circuits of the NAc with pathway- and cell-specific plasticity. Together these data reveal a high degree of synapse and circuit specificity in the adult NAc core.

## MATERIALS AND METHODS

### Animals

Animals were treated in compliance with the European Communities Council Directive (86/609/EEC) and the United States National Institutes of Health Guide for the care and use of laboratory animals. The French Ethical committee authorized this project (APAFIS#3279-2015121715284829 v5). Male Drd1a-tdTomato mice were from The Jackson Laboratory (Bar Harbor, ME, USA) and female C57Bl/6J background mice were purchased from the Janvier Laboratory (Le Genest-Saint-Isle, France). Mice were acclimated to the animal facility for one week and then housed in male and female pairs to enable breeding of hemizygous offspring. Mice were ear punched for identification and genotyping. Mice were housed at constant room temperature (20 ± 1°C) and humidity (60%) and exposed to a light cycle of 12h light/dark (8:00 a.m. to 08:00 p.m.), with ad libitum access to food and water. All synaptic plasticity experiments were performed on male offspring C57BL/6J mice between P90 and P130.

### Injection of the virus

Microinjection needles (32G) were connected to a 10 µL Hamilton syringe and filled with purified, concentrated adeno-associated virus (1.98×1013 infectious units per mL) encoding hChR2-EYFP under control of the CaMKIIα promoter (University of Pennsylvania, Philadelphia, USA). Mice were anesthetized with 150 mg/kg ketamine and 50 mg/kg xylazine and placed in a stereotaxic frame. Microinjection needles were bilaterally placed into the vHipp (Coordinates: AP=-3.2mm; ML=2.85mm; DV=3.84mm), basolateral amygdala (Coordinates: AP=-1mm; ML=+3.1mm; DV=3.9mm) or prefrontal cortex (Coordinates: AP=2mm; ML=0.3mm; DV=2mm) and 250 nL virus was injected with a speed of 100 nL/min. The needles were left in place for an additional 5 min to allow for diffusion of virus particles away from injection site.

### Slice preparation

Five to six weeks after surgery, adult male mice (P100-P130) were deeply anesthetized with isoflurane and sacrificed as previously described (Robbe et al., 2002; Lafourcade et al., 2011; Jung et al., 2012; Neuhofer et al., 2018). The brain was sliced (300 μm) on the coronal plane with a vibratome (Integraslice, Campden Instruments) in a sucrose-based solution at 4°C (in mm as follows: 87 NaCl, 75 sucrose, 25 glucose, 2.5 KCl, 4 MgCl_2_, 0.5 CaCl_2_, 23 NaHCO_3_ and 1.25 NaH_2_PO_4_). Immediately after cutting, slices containing the NAc were stored for 1 h at 32°C in a low calcium ACSF that contained (in mm) as follows: 130 NaCl, 11 glucose, 2.5 KCl, 2.4 MgCl_2_, 1.2 CaCl_2_, 23 NaHCO_3_, 1.2 NaH2PO_4_, and were equilibrated with 95% O2/5% CO2 and then at room temperature until the time of recording. EYFP expression was examined in slices containing the virus injection sites to assess placement accuracy.

### Electrophysiology

Whole-cell patch-clamp recordings of visualized NAc medium spiny neurons (MSNs) were made in coronal slices containing the NAc as previously described (Robbe et al., 2002; Lafourcade et al., 2011; Jung et al., 2012; Neuhofer et al., 2018). During the recording, slices were placed in the recording chamber and superfused at 2 mL/min with normal ACSF. All experiments were done at 25°C. The superfusion medium contained picrotoxin (100 μM) to block GABA Type A (GABA-A) receptors, unless specified otherwise. All drugs were added at the final concentration to the superfusion medium.

For whole-cell patch-clamp experiments, neurons were visualized using an upright microscope with infrared illumination. The intracellular solution was based on K+ gluconate (in mM as follows: 145 K+ gluconate, 3 NaCl, 1 MgCl_2_, 1 EGTA, 0.3 CaCl_2_, 2 Na^+^ATP, and 0.3 Na^+^GTP, 0.2 cAMP, buffered with 10 HEPES). The pH was adjusted to 7.2 and osmolarity to 290–300 mOsm. Electrode resistance was 4–6 MOhm.

A −2 mV hyperpolarizing pulse was applied before each evoked EPSC to evaluate the access resistance, and those experiments in which this parameter changed >25% were rejected. Access resistance compensation was not used, and acceptable access resistance was <30 MOhm. The potential reference of the amplifier was adjusted to zero before breaking into the cell. Cells were held at −70 mV.

Current-voltage (I–V) curves were made by a series of hyperpolarizing to depolarizing current steps immediately after breaking into the cell. Membrane resistance was estimated from the I– V curve around resting membrane potential (Kasanetz et al., 2010; Thomazeau et al., 2014; Martin et al., 2017).

The paired-pulse ratio (PPR) was calculated from the peak current of two closely spaced EPSCs (50 ms), such that the PPR = Peak 2/Peak 1. Quoted PPR values are the average of 30 trials. For the measurements of quantal EPSCs (qEPSCs), transmitter release was desynchronized by substituting calcium with strontium (4 mM) in the superfused ACSF. Asynchronous EPSCs were examined during a 200 ms window beginning 5 ms after optical stimulation. Recordings were analyzed if the frequency of events in this 200 ms window were significantly greater than during the 200 ms window preceding the stimulation as previously described (Britt et al., 2012).

### Optogenetics

A 473 nm laser (Dragon Laser, Changchun Jilin, China) coupled to a 50 μm core glass silica optical fiber (ThorLabs) was positioned directly above the slice orientated at 30° approximately 350 μm from the recording electrode. At the site of recording discounting scattering a region of approximately 0.05 mm2 was illuminated that after power attenuation due to adsorption and scattering in the tissue was calculated as approximately 100 mW/mm^2^ (Yizhar et al., 2011). Optically evoked EPSCs were obtained every 10 s with pulses of 473 nm wavelength light (0-10 mW, 2 ms).

### Drugs

Drugs were added at the final concentration to the recording ACSF media. Picrotoxin, Strontium and Tetrodotoxin from Sigma (St. Quentin Fallavier, France); Capsazepine and CP55,940 from Tocris Cookson (Bristol Bioscience, UK); CNQX from the National Institute of Mental Health’s Chemical Synthesis and Drug Supply Program (Rockville, MD, USA); NESS 0327 from Cayman Chemical (Bertin Bioreagent, St. Quentin en Yvelines, France).

### Data Acquisition and Analysis

Whole-cell patch-clamp recordings were performed with an Axopatch-200B amplifier as previously described (Robbe et al., 2002; Kasanetz and Manzoni, 2009; Lafourcade et al., 2011; Jung et al., 2012; Thomazeau et al., 2014, 2017; Martin et al., 2017). Data were low pass filtered at 2 kHz, digitized (10 kHz, DigiData 1440A, Axon Instruments), collected using Clampex 10.2 and analyzed using Clampfit 10.2 (all from Molecular Device). The magnitude of plasticity was calculated and compared 25-30 min after induction.

Spontaneous and quantal AMPAR-mediated EPSCs (sEPSCs/qEPSCs) were recorded using Axoscope 10 (Molecular Devices). sEPSCs/qEPSCs were filtered at 2 kHz and digitized at 20 kHz. sEPSCs/qEPSCs amplitude and frequency were analyzed with Axograph X using a double exponential template: f(t)=exp(−t/rise)+exp(−t/decay), rise=0.5 ms. The threshold of amplitude detection was set at 5 pA.

Statistical analysis of data was performed with Prism (GraphPad Software 6.0) using tests indicated in the main text after outlier subtraction (ROUT test). N values represent cells or individual animals. All values are given as mean ± SEM, and statistical significance was set at *p < 0.05.

## RESULTS

We recorded a total of 412 medium spiny neurons (MSNs) from the NAc core of 291 Tg (Drd1a-tdTomato) Calak hemizygous adult male mice (P100-130). MSNs were either tdTomato-labeled “D1a-positive” MSNs or tdTomato-unlabeled/presumably “D1a-negative” MSNs. Because previous studies have consistently shown that unlabeled D1a negative MSNs are all D2 positive MSNs, in the remainder of the study we refer to tdTomato-unlabeled MSNs as D2 MSNs (Bertran-Gonzalez et al., 2008; Ade et al., 2011; Enoksson et al., 2012; Thibault et al., 2013; Cao et al., 2018).

### Intrinsic properties of D1 and D2 medium spiny neurons in adult Drd1a-tdTomato mice

In juvenile and adolescent mice, D2 MSNS are more excitable than D1 MSNs in the NAc (Grueter et al., 2010; Ma et al., 2014; Cao et al., 2018). To our knowledge, the intrinsic properties of D1 and D2 MSNs in adult NAc, *a fortiori* in the NAc core, have not been described. The intrinsic properties of current-clamped and visually identified neighboring D1 and D2 MSNs were compared in NAc core slices from adult *Drd1a-td*Tomato mice (Figure 1). The membrane reaction profiles of D1 and D2 MSNs in response to a series of somatic current steps differ greatly (Figure 1B). Differences between MSN subtypes extended to their resting membrane potential which was significantly more depolarized in D1-than in D2 MSNs (Figure 1C) and the rheobase that was significantly lower in D1-than D2 MSNs (Figure 1D). The “hyper-excitability” of D1 MSNs was accompanied by an increased number of action potentials in response to somatic currents steps in D1-compared to D2 MSNs (Figure 1E). Action potential duration was also shorter in D1 MSNs and action potential after-hyperpolarization (fAHP) was larger in D2 MSNs (Figure 1-1). Thus, in the NAc core of adult mice, D1 MSNs are more excitable than D2 MSNs.

**Figure 1:**
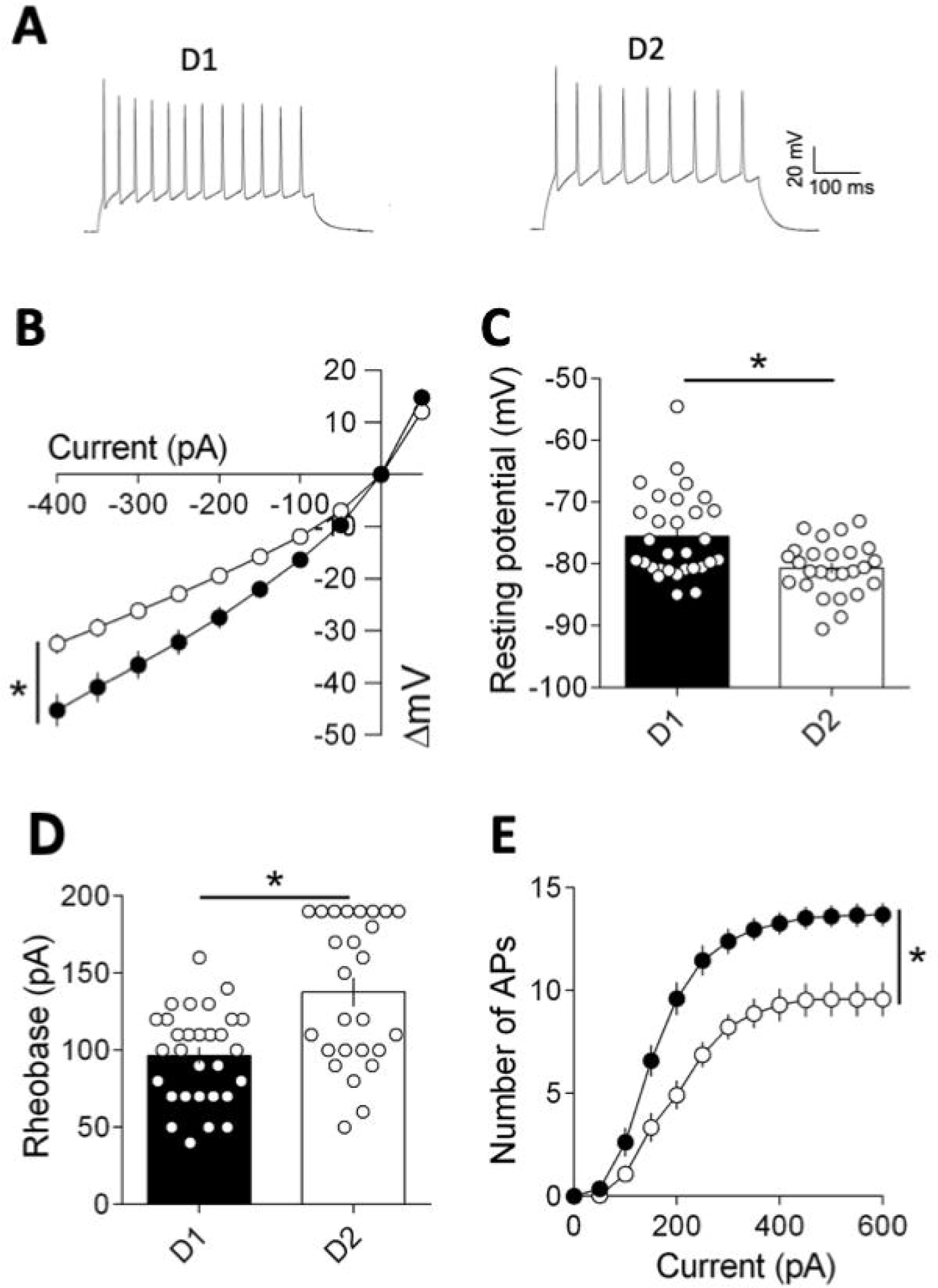
Adult NAc core D1 MSNs are more excitable than D2 MSNs. **A)** Typical membrane responses from NAc core D1 or D2 MSNs in reaction to a series of increasing somatic current injections. Sample spike trains in response to depolarizing current from D1 or D2 MSN. **B)** Summary of all the current–voltage (I–V) curves recorded in D1 (black, n=30) and D2 (white, n=26) MSNs (F_(interaction 26,1404)_=8.03, p<0.0001, F_(cell type 1,54)_=2.893, p<0.0001, two-way repeated measures ANOVA). **C)** The resting membrane potential of D2 MSNS was significantly hyperpolarized compared to that of D1 MSNs (p=0.0051, Mann-Whitney U test). **D)** The rheobase, the minimal current required to trigger an action potential, was much lower in D1 MSNs (p=0.0025, Mann-Whitney U test). **E)** The number of evoked action potentials in response to increasing depolarizing current steps was larger in D1 MSNs compared to D2 MSNs (F_(interaction 12,648)_= 5.927, p<0.0001, F_(cell type 1,54)_=27.38, p<0.0001, two-way repeated measure ANOVA). Individual point in scatter dot plots represents one individual neuron. All values are represented as mean ± SEM. *p<0.05.

### Cell-type specific hierarchy of excitatory inputs in the adult NAc core

In postsynaptic NAc shell MSNs, optogenetic methods and targeted channelrhodopsin-2 (ChR2) expression to projection neurons from ventral hippocampus (vHipp), the basolateral amygdala (BLA) and the prefrontal cortex (PFC) revealed that vHipp inputs elicit the largest excitatory currents (vHipp>BLA>PFC; Britt et al., 2012). At the strict anatomical level, the PFC preferentially projects to the NAc core, in contrast with the vHipp which preferentially projects to the shell. The BLA projects equally to both NAc core and shell (Li et al., 2018).

However, the functional hierarchy of specific excitatory synaptic inputs to identified MSN subtypes in the NAc core is largely unknown. Akin to Britt and collaborators (Britt et al., 2012), we targeted ChR2 expression to projection neurons in the VHipp, BLA and PFC in order to compare the functional strength and synaptic properties of these inputs onto identified D1 and D2 MSNs in the NAc core of adult mice (Figure 2 and 3; Figure 2-1).

**Figure 2:**
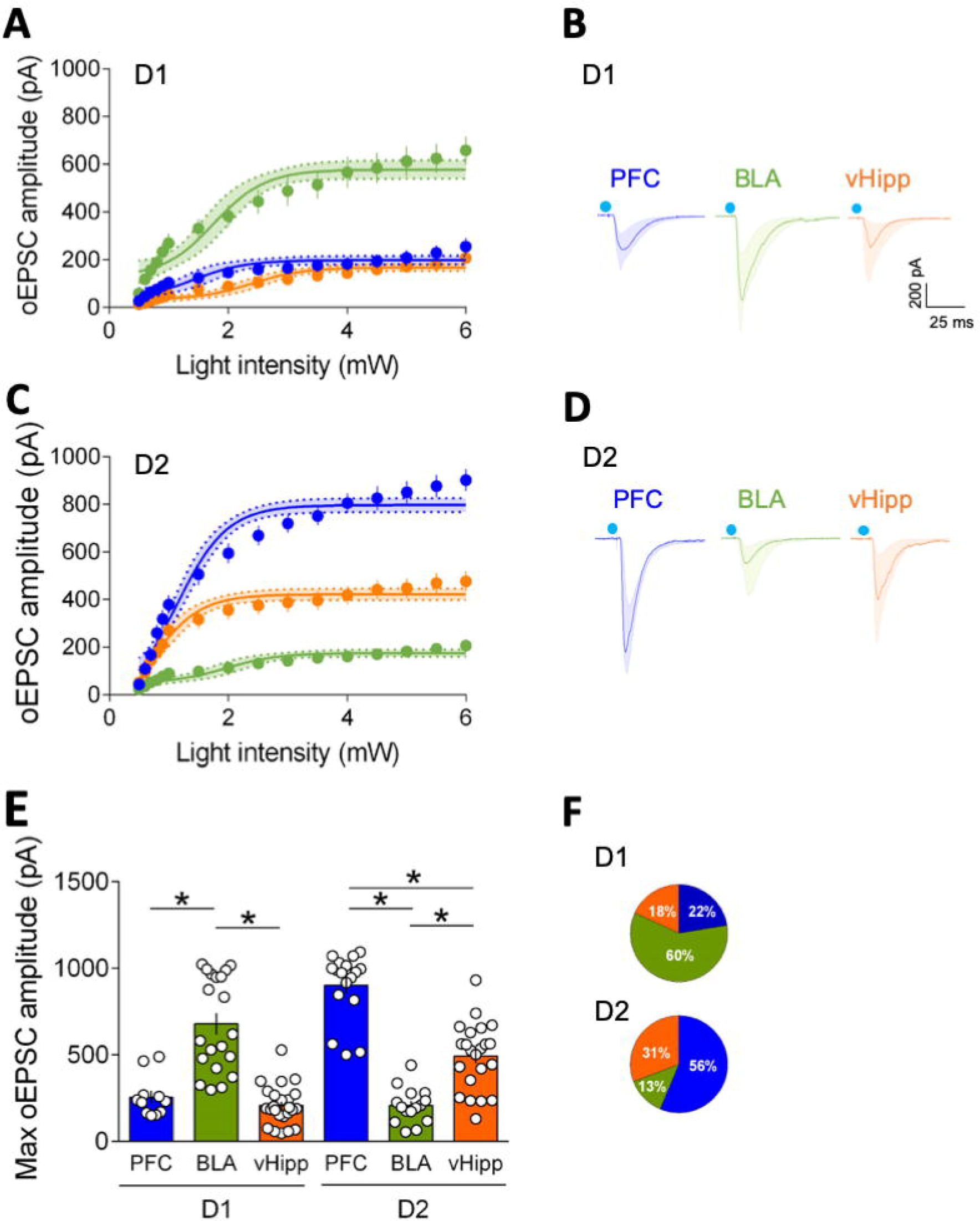
The hierarchy of optically driven synaptic inputs depends on the cellular identity of the target MSNs. **A)** In D1 MSNs the largest excitatory oEPSCs were elicited at BLA fibers (green, n=20) followed by PFC (blue, n=11) and vHipp (orange, n=24). (F_(light intensity × input 30,832)_=5.305, p<0001, F_(input 2,832)_=359.4, p<0001, two-way repeated measures ANOVA, 95% Confidence Interval). B) Example oEPSC traces for PFC, BLA and vHipp inputs in D1 MSNs. **C)** In D2 MSNs, the largest oEPSCs were elicited by PFC inputs (n=17) followed by vHipp (n=22) and the BLA (n=15). (F_(light intensity × input 30,816)_=11.12, p<0001, F_(input 2,816)_= 543.8, p<0001, two-way repeated measures ANOVA, 95% Confidence Interval). **D)** Example oEPSC traces for PFC, BLA and vHipp inputs in D2 MSNs. E) Summary bar histogram of maximum oEPSCs for PFC, BLA and vHipp inputs in D1 and D2 MSNs (F_(input × cell type 2,103)_=78.33, F_(cell type 1,103)_=18.00, p<0001, F_(input 2,103)_=13.41, p<0001, two-way ANOVA). **F)** Circle plots showing relative distribution of oEPSCs responses for PFC, BLA and vHipp inputs in D1 and D2 MSNs. Individual point in scatter dot plots represents one individual neuron. All values are represented as mean ± SEM or geometric mean ± CI. *p<0.05.

**Figure 3:**
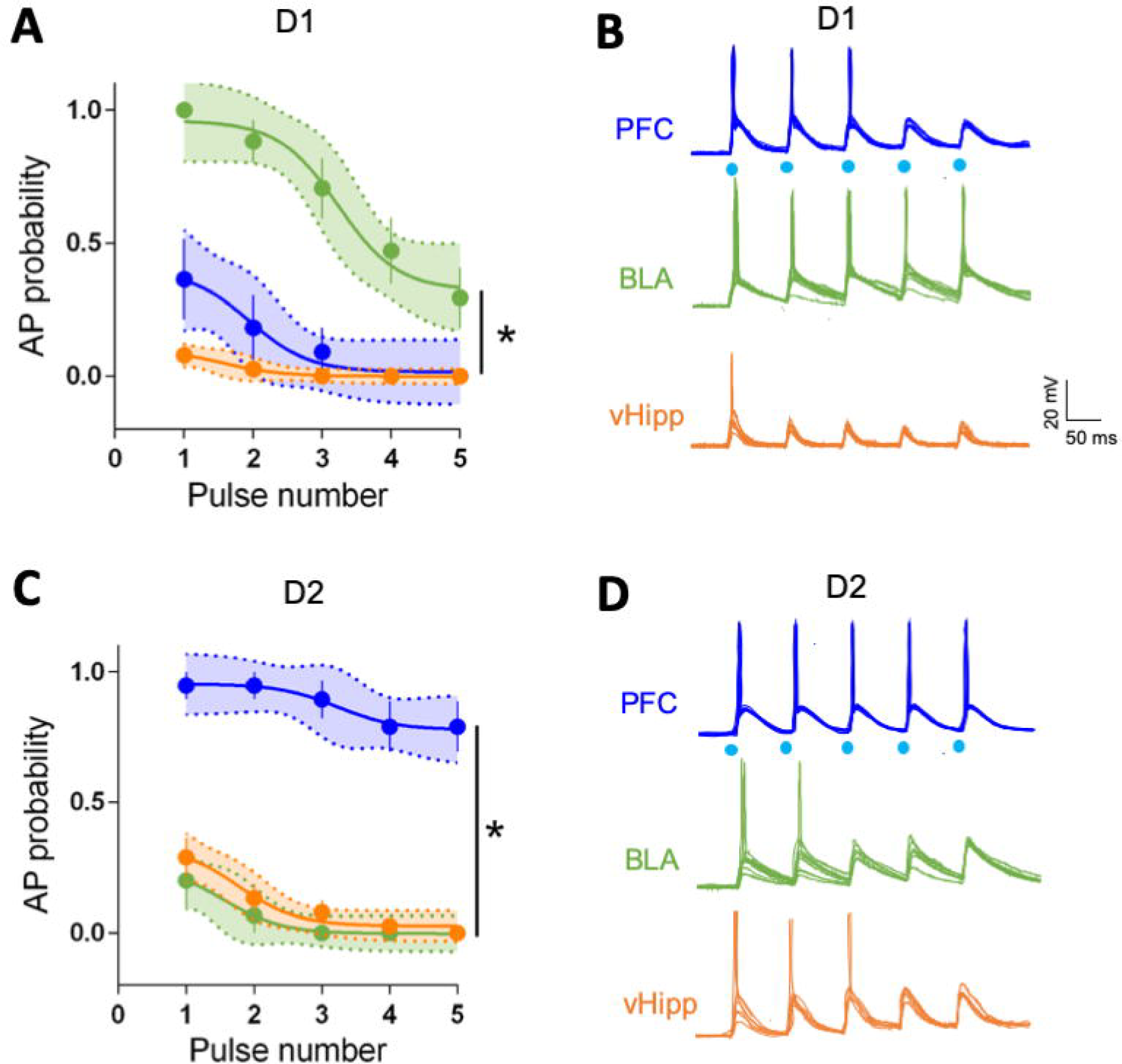
Pathway-specific drive of action potential in identified NAc core MSNs. **A)** Comparison of the probability of AP firing versus pulse number shows that BLA (green) inputs preferentially drive the firing of D1 MSNs (PFC n=6, BLA n=7, vHipp n=6, F_(pulse number × AP probability 8,315)_=4.505, p<0001, F_(input 2,315)_=148.9, p<0001, two-way repeated measures ANOVA, 95% Confidence Interval). **B)** Example traces of APs evoked in D1 MSNs, in response to trains of optical stimulation of PFC, BLA and vHipp inputs (2 ms light pulses, 5 pulses at 10 Hz). **C)** Comparison of the probability of AP firing versus pulse number shows that PFC (blue) inputs preferentially drive the firing of D2 MSNs (PFC n=6, BLA n=6, vHipp n=6, F_(pulse number × AP probability 8,345)_= 0.3976, p=0.921605, F_(input 2,345)_=256,6, p<0001, two-way repeated measures ANOVA, 95% Confidence Interval). **D)** Example traces of APs evoked in D2 MSNs in response to trains of optical stimulation of PFC, BLA and vHipp inputs (2 ms light pulses, 5 pulses at 10 Hz). Blue dots indicate time of stimulation. All experiments performed in the presence of picrotoxin to prevent from feedforward inhibition. All values are represented as mean ± SEM or geometric mean ± CI. *p<0.05.

Five to six weeks after viral infection with pAAV9-CaMKIIa-hChR2(H134R)-EYFP into the vHipp, BLA or PFC, strong expression of ChR2-EYFP was observed in the NAc core (Figure 2-1A). To best mimic “real life” action potentials and avoid direct illumination of axon terminals, light pulses (473nm) were delivered with a custom-made optical fiber placed approximately 350 µm from the recorded MSNs (Figure 2-1B). Independently of input, light-evoked excitatory postsynaptic currents (“optical EPSCs”, oEPSCs) were abolished in the presence of the ionotropic glutamate receptor antagonist CNQX (20 µM) or the sodium channel blocker tetrodotoxin (TTX, 1 µM) (Figure 2-1C). These control experiments show that oEPSCs genuinely are glutamate-mediated EPSCs induced following the light-evoked activation of axonal Na^+^ channels.

In the NAc shell, “irrespective of which pathway was optically stimulated, oEPSCs were observed in more than 95% of recorded neurons” (Britt et al., 2012). In our experiments the same held true (data not shown), irrespective of MSN subtypes suggesting that each MSN receives innervation from PFC, vHipp and BLA (Finch, 1996; Groenewegen et al., 1999; French and Totterdell, 2002, 2003; McGinty and Grace, 2009; Britt et al., 2012) (Figure 2-2).

Using increasing optical stimulations and whole-cell patch clamp of visually identified MSNs, we built and compared input-output curves for PFC, BLA and vHipp inputs at both D1 and D2 MSNS, in the presence of picrotoxin (100 µM) to abolish feedforward inhibition (Figure 2, see also Figure 5). Evoked oEPSCs exhibited a consistent profile distribution in response to increasing stimulation intensity across different pathways (Figure 2A). We found that, in D1 MSNs, the largest excitatory currents were elicited by optical stimulation of BLA fibers followed by PFC and vHipp (D1 MSNs: BLA>PFC=vHipp; Figure 2A, B, E, F). In D2 MSNs, the largest oEPSCs were elicited by PFC inputs followed by vHipp; the BLA sent the weakest projections to D2 MSNs in marked contrast to D1 MSNs (D2 MSNs: PFC>vHipp>BLA; Figure 2C, D, E, F). These data show that in the adult NAc core, the hierarchy of synaptic inputs is specific to the cellular identity of the target MSNs.

### Input- and cell-type-specific drive of action potential firing in the adult NAc core

What is the relationship between maximal excitatory synaptic strength and action potential firing of identified MSNs? We first sought to determine if the pathway/cell specificity extended to the ability of different pathways to drive postsynaptic action potentials in identified MSNs. Current-clamp recordings of adult D1 MSNs revealed that optical recruitment of BLA inputs elicited postsynaptic action potential, with greater probability than either PFC or vHipp inputs (Figure 3A, B). In marked contrast, PFC inputs onto D2 MSNs were the most likely to trigger action potentials, followed by BLA and vHipp (Figure 3C, D). Performed in the absence of feedforward inhibition, i.e. in the presence of picrotoxin, these results faithfully mirror the aforementioned cell-type specificity of the hierarchy of synaptic strength (Figure 2).

### Feedforward inhibition in the adult NAc core is input- and cell-type-specific

Next, we decided to compare the weight of feedforward inhibition across identified microcircuits (Figure 4). To this aim we measured spontaneous and evoked firing of identified MSNs in the absence and presence of the GABA-A receptor antagonist picrotoxin (Figure 4). The consequences on spontaneous firing of blocking GABA-A receptors greatly differed in cell-attached D1 and D2 MSNs (Figure 4A). Spontaneous firing increased in both D1 and D2 MSNs, but disinhibition was larger in D1 than D2 MSNs, suggesting that continuous inhibitory bombardment is high in D1. To evaluate feedforward inhibition, we compared the effects of picrotoxin on the spike ratio during a two second window before and after opto-stimulation. Blocking feedforward inhibition had minimal effects at hippocampal inputs onto both cell types (Figure 4B, **right**), while it doubled the firing rate recorded in D1 and D2 MSNs in response to BLA stimulation (Figure 4B, **middle**). At PFC inputs, feedforward inhibition was cell-type specific: it was minimal at D1 and maximal at D2 MSNs. These data reveal a previously undiscovered high level of pathway/cell specificity in feedforward inhibition in NAc core MSNs.

**Figure 4:**
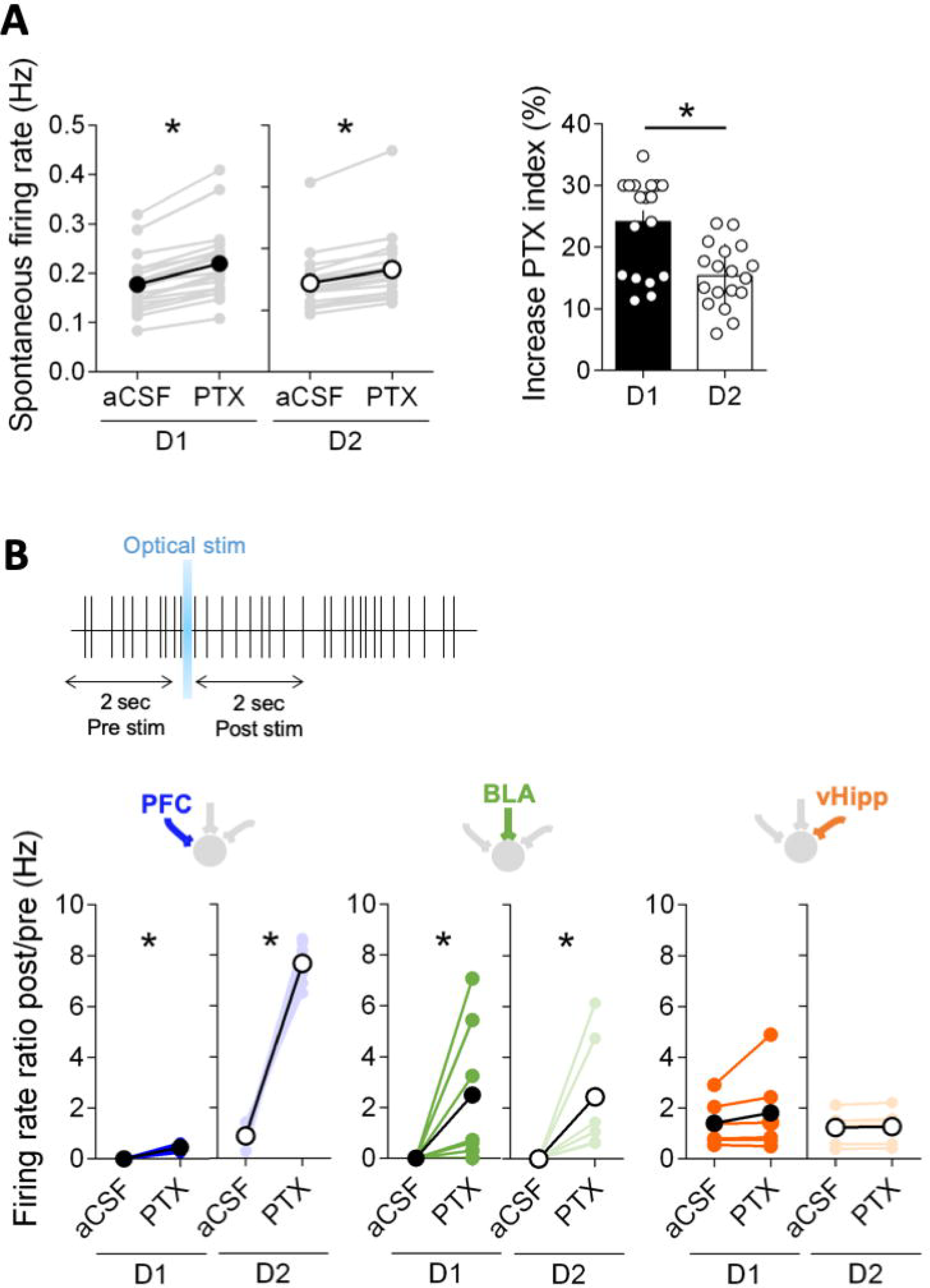
Pathway and cell-type specific feedforward inhibition in NAc core MSNs. **A)** Basal synaptic inhibition is larger in D1 (black, n=19) than D2 MSNs (white, n=18). Bath application of the GABA-A receptors antagonist picrotoxin increased the spontaneous firing of both cell-attached D1 and D2 MSNs (left panel) p<0001, paired t-test) but to a different extent: the disinhibition was larger in D1 than D2 MSNs, suggesting different basal levels of synaptic inhibition (right panel) (p=0.0013, Mann-Whitney U test). **B)** Comparing firing rate ratio during a two second window before and after pathway specific opto-stimulation in the absence (aCSF) and presence of picrotoxin (PTX) reveals that feedforward inhibition is pathway specific. At PFC inputs (blue), feedforward inhibition was cell specific. Blocking GABA-A receptors had almost no effect on the PFC-driven firing of D1 MSNs (n=6) but yielded to a large increase of PFC-driven firing of D2 (n=6). Blocking feedforward inhibition doubled the firing rate of both D1 and D2 MSNs in response to BLA stimulation (green, D1 MSNs n=7, D2 n=6), but had minimal effects at hippocampal inputs onto both cell types (orange, D1 MSNs n=6, D2 n=6). (PFC: D1 MSNs, p=0.0007 and D2 MSNs p<0.0001; BLA: D1 MSNs, p<0.0001 and D2 MSNs, p<0.0001; vHipp: D1 MSNs, p=0.2607 and D2 MSNs, p=0.0919, paired t-test). Individual point in scatter dot plots represents one individual neuron. All values are represented as mean ± SEM. *p<0.05.

### Input- and cell-type-specific synaptic properties in the adult NAc core

To investigate cell-type specific connectivity at glutamatergic inputs into the NAc, we compared post- and pre-synaptic parameters at disambiguated synapses in the NAc core. First, we measured the paired-pulse ratio (PPR), a classical index of neurotransmitter release probability (Silver et al., 1998), Figure 5A,B). In D1 MSNs, BLA inputs had a low PPR/high release probability and the highest PPR/lowest release probability was found at PFC inputs (D1: BLA>vHipp>PFC). In contrast, in D2 MSNs, PFC and BLA inputs exhibited the highest and lowest release probability, respectively, while the lowest probability of release was found at BLA inputs (D2: PFC=vHipp>BLA).

**Figure 5:**
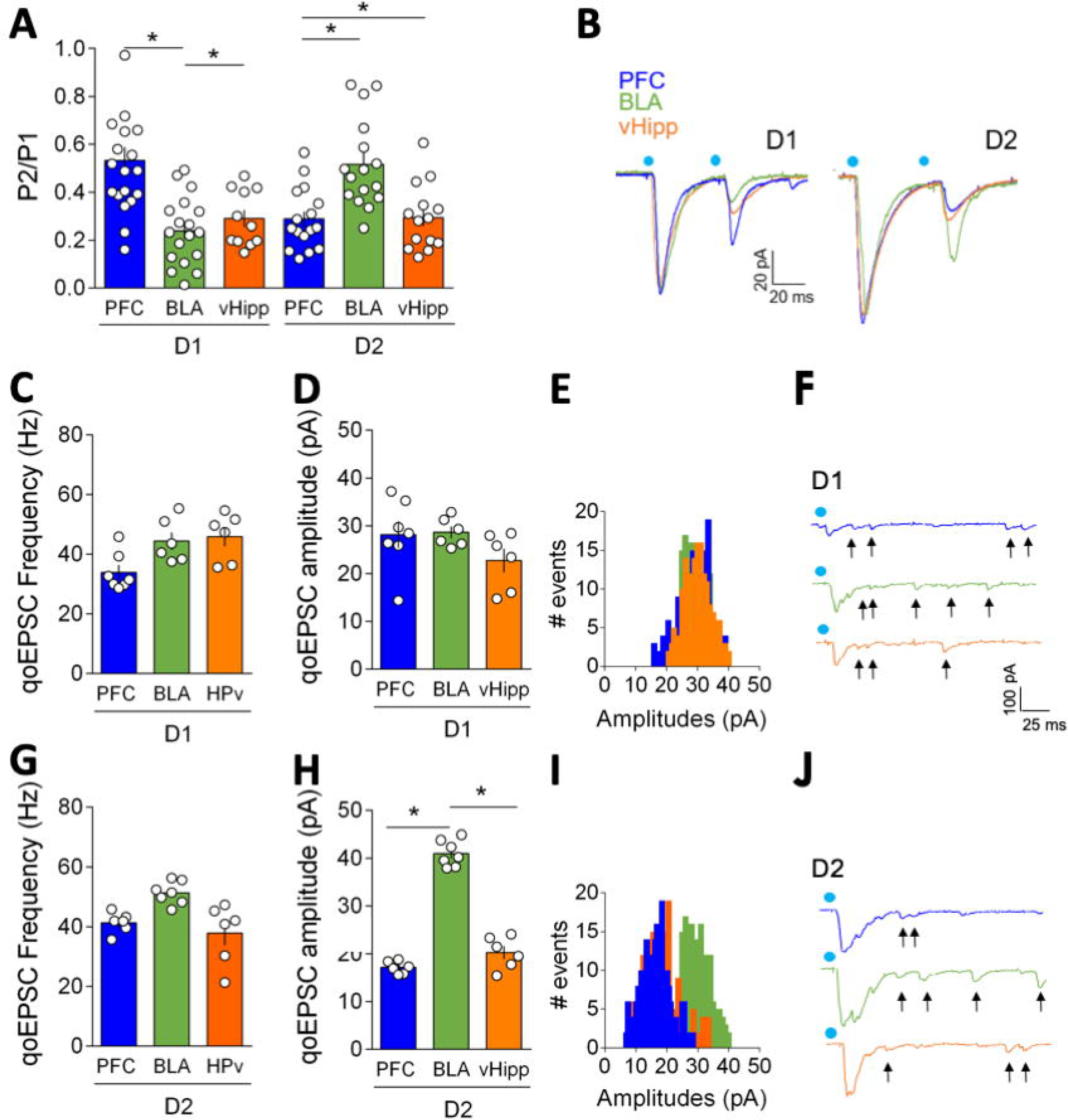
Input- and cell-type-specific post and presynaptic properties in the NAc core. **A)** Average paired-pulse ratio (P2/P1) measured in D1 and D2 MSNs (voltage-clamp,-70mV) in response to paired optical stimulations (50msec interval) of vHipp (orange), BLA (green) and PFC (grey) inputs (D1 MSNs: PFC n=12, BLA n=16, vHipp n=15; D2 MSNs: PFC n=19, BLA n=18, vHipp n=11, F_(input × cell type 2,85)_=16.48, p<0.0001, F_(input 2,85)_=4.847, p=0.0102, two-way repeated measure ANOVA). In D1 MSNs, PFC inputs had a high PPR/low release probability compared to BLA and vHipp inputs that both exhibited a low PPR/high release probability (BLA>vHipp>PFC). In D2 MSNs, the lowest release probability was observed at BLA inputs compared to PFC and vHipp inputs (PFC=vHipp>BLA). **B)** Example traces of evoked paired oEPSC responses in D1 (left) and D2 (right) MSNs. **C-J** Light-evoked input-specific quantal EPSCs (qoEPSC) in the adult NAc core. In D1 MSNs qoEPSCs frequency **(C)** and qoEPSCs amplitude **(D)** were similar across pathways (PFC n=7, BLA n=6, vHipp n=6). **E)** Histograms for representative D1 cells showing the distribution of qoEPSC amplitude across all trials. **F)** Representative qoEPSCs recorded in D1 MSNs in response to pathway specific opto-stimulation (arrows indicate detected qoEPSCs). In D2 MSNs **G)** qoEPSC frequencies were similar at PFC, BLA and vHipp inputs. **H)** qoEPSCs’ amplitudes were larger at BLA inputs compared to PFC and vHipp afferents (PFC n=6, BLA n=7, vHipp n=6, F_(input 2,16)_=162.6, p<0.0001, one-way ANOVA). **I)** Histograms for representative D2 cells showing the distribution of qoEPSC across all trials. **J)** Representative qoEPSCs recorded in D2 MSNs in response to pathway specific opto-stimulation (arrows indicate detected qoEPSCs). Blue dots indicate optical stimulations. Individual point in scatter dot plots represents one individual neuron. All values are represented as mean ± SEM. *p<0.05.

In immature mice, miniature EPSCs properties were similar in D1 and D2 MSNs in early life (P16-24, Cao et al., 2018), while around P42 spontaneous EPSCs (sEPSC) were more frequent in D2 MSNs (Ma et al., 2012) in disagreement with an earlier report of higher miniature EPSCs in D1 MSNs (P28-56, Grueter et al., 2010). We recorded sEPSC in D1 and D2 NAc core MSNs of adult mice (Figure 5-1). The amplitude of the sEPSCs measured in D1 MSNs was slightly larger than that measured in D2 MSNs (in agreement with Grueter et al., 2010). Both MSN subtypes exhibited similar distribution and average inter-event intervals.

Next, we measured light-evoked input-specific quantal EPSCs (qoEPSCs) by replacing our intracellular medium’s calcium with strontium to desynchronize transmitter release (Figure 5) (Britt et al., 2012; McGarry and Carter, 2017). qoEPSCs amplitude provide a direct estimate of postsynaptic efficacy measure and qoEPSC frequency was taken as an indirect indication of the number of connections (Goda and Stevens, 1994). In D1 MSNs, qoEPSC’s amplitude and frequency were similar across pathways and cell types (Figure 5C-E). In contrast, qoEPSC’s amplitude and frequency were larger at BLA-D2 synapses compared to the other afferents. These findings are compatible with the idea that BLA afferents make more efficacious and/or numerous synapses onto D2-than to D1 MSNs.

### Pathway specific expression of CB1R and TRPV1R endocannabinoid receptors

The endogenous cannabinoid (eCB) system modulates synaptic circuits in the CNS and notably the striatal areas (Araque et al., 2017). Inhibitory CB1 receptors (CB1R) have long been described at glutamatergic NAc core synapses using standard electrical stimulation (Robbe et al., 2001). Stimulation of CB1R with a submaximal dose of a synthetic cannabimimetic (CP55940, Figure 6A, B) inhibited oEPSC in D1 and D2 MSNs, irrespectively of the input pathway, in support of the idea that functional CB1R are present at all of these inputs. The degree of inhibition induced by the single dose used here, however, was pathway dependent. Thus, CP55940-induced inhibition was larger at PFC and vHipp-D1 synapses (compared to BLA, Figure 6C) and vHipp-D2 synapses (compared to PFC and BLA, Figure 6D).

**Figure 6:**
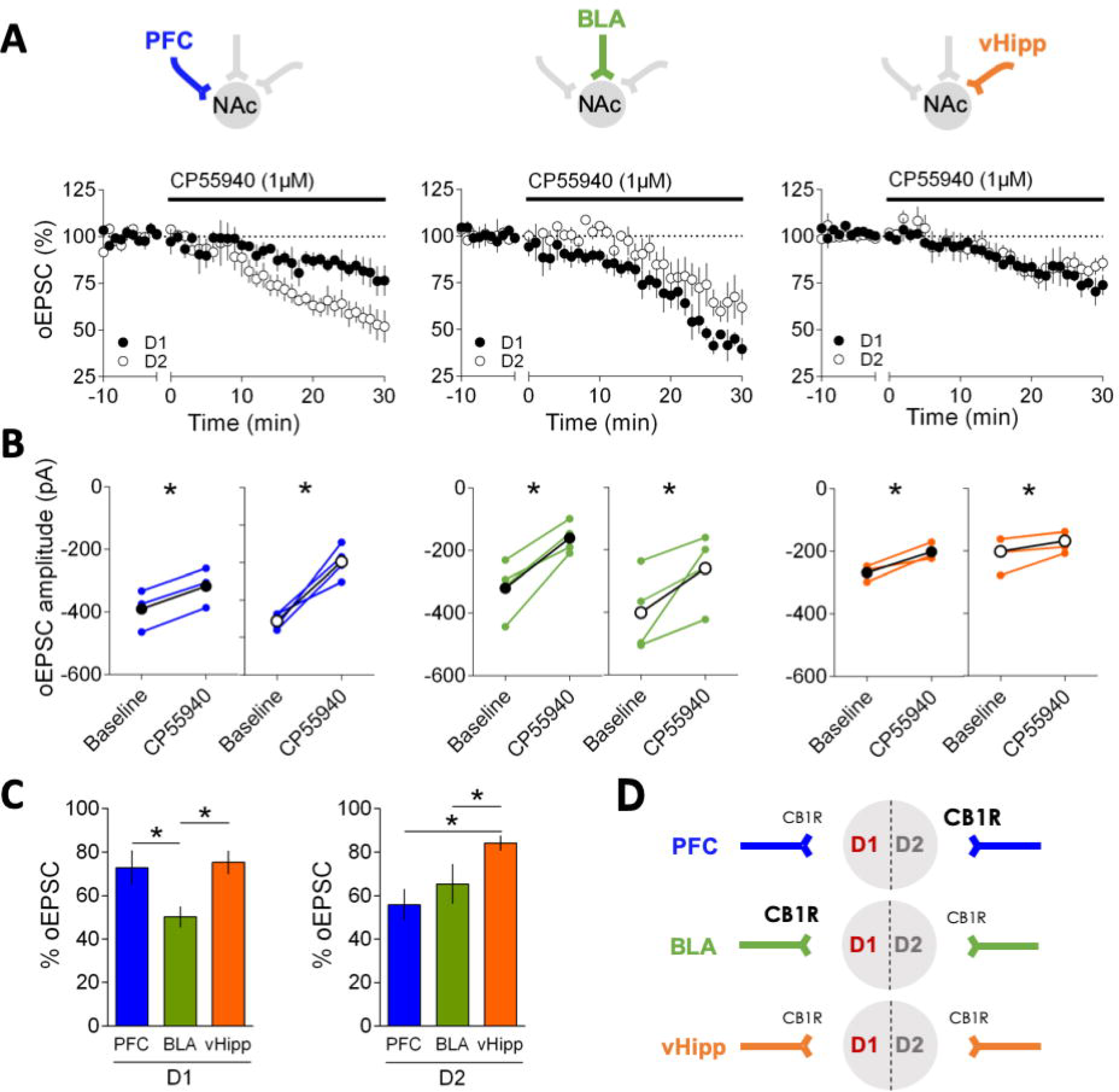
CB1R inhibition is common to all inputs onto D1 and D2 NAc core MSNs. **A)** Bath application of the synthetic CB1R agonist CP55940 (1 µM), inhibited oEPSC in D1 (black) and D2 (white) MSNs. CP55940 was applied for 30 min (black bar) after at least 10 min of stable baseline recording. **B)** Individual and averaged oEPSCs amplitude experiments before (baseline) and 25-30 min after CP55940. (D1 MSNs, black circles: PFC n= 3, p=0.0014, BLA n=4, p=0.0110, vHipp n=3, p=0.0411; D2 MSNs, white circles: PFC n=4, p=0.0114, BLA n=4, p=0.0188, vHipp n=4, p=0.0496 paired t-test). **C)** Summary bar histogram comparing CP55940-induced inhibition of oEPSCs at identified PFC, BLA and vHipp inputs onto D1 (PFC n= 3, BLA n=4, vHipp n=3, F_(input 2,7)_=14.32, p=0.0034, one-way ANOVA) and D2 MSNs (PFC n=4, BLA n=4, vHipp n=4, F_(input 2,9)_=4.207, p=0.0513, one-way ANOVA). **D)** Schematic view of the relative weight of CB1R-mediated inhibition at PFC, BLA and vHipp inputs onto D1 and D2 MSNs. n represents the number of mice. All values are represented as mean ± SEM. * p<0.05.

The nonselective cation channel, transient receptor potential cation channel subfamily V member 1 (TRPV1), is a multifaceted mediator of eCB signaling in the CNS (Gibson et al., 2008; Manduca et al., 2017; Bara et al., 2018), notably in the striatum: pharmacological activation of TRPV1R suppresses and facilitates transmitter release in the dorsal striatum (Musella et al., 2009) and inhibits excitatory inputs onto D2 NAc core MSNs (Grueter et al., 2010). Multiple inputs onto D1 and D2 MSNs differentially expressing various levels of TRPV1R could explain these apparently contradicting results. Thus, we compared the effects of the specific TRPV1R agonist capsaicin in the disambiguated NAc core synapse preparation (Figure 7). While capsaicin inhibited PFC-evoked oEPSCs in both D1 and D2 MSNs, the TRPV1R agonist had many-sided effects on the other pathways. Capsaicin had opposite effects on BLA-evoked oEPSCs depending on the MSNS subtypes: the efficacy of BLA D1 MSNs synapses was enhanced in the presence of the TRPV1 agonist while that at BLA-D2 MSNs synapses was reduced. Finally, at vHipp inputs, TRPV1R activation specifically inhibited vHipp-D1 synapses and had no effect at vHipp synapses onto D2 MSNs.

**Figure 7:**
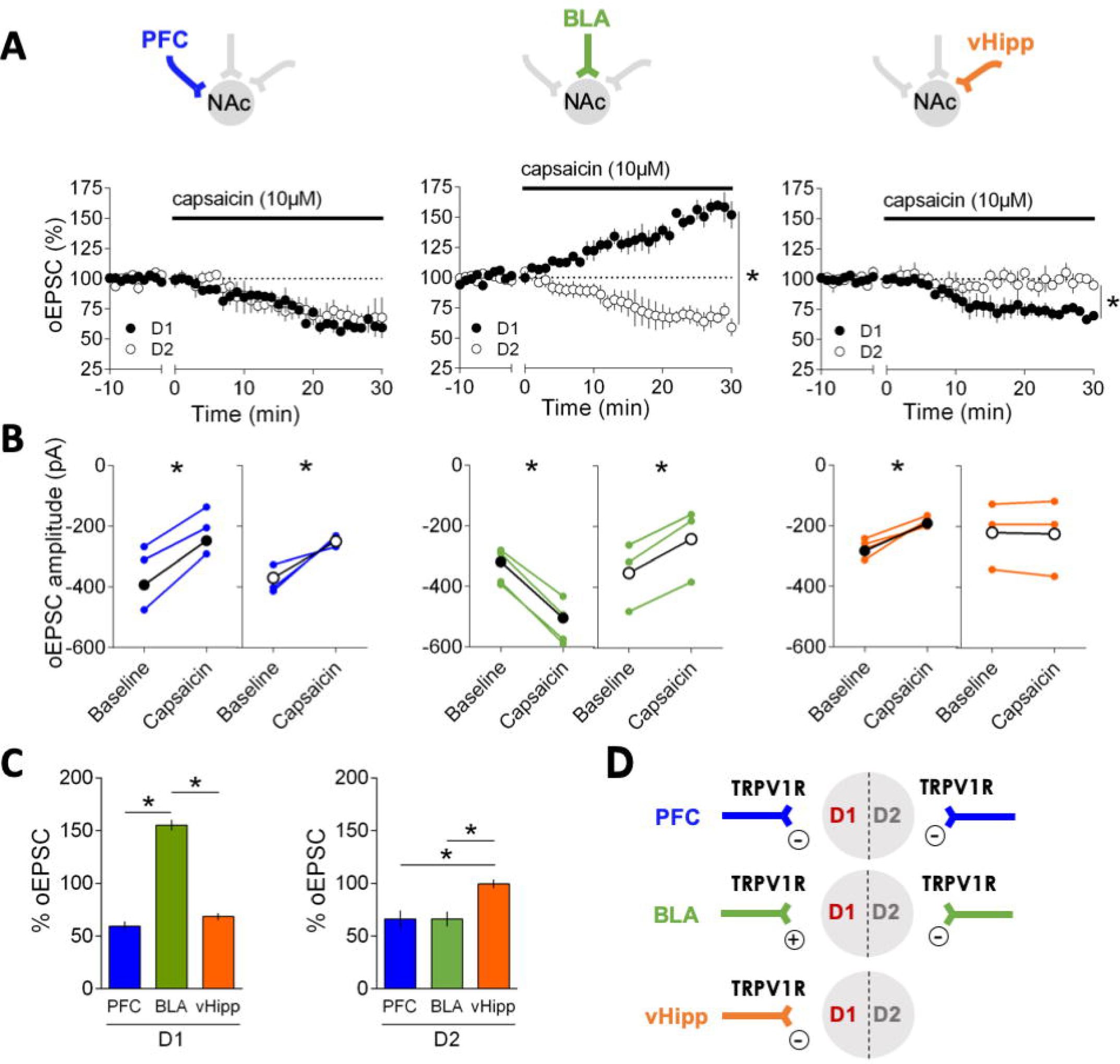
Divergent input and cell specific control of excitatory NAc synapses by TRPV1R. **A)** Effects of the TRPV1R agonist capsaicin (10µM) on oEPSCs at PFC, BLA and vHipp synapses onto D1 (black circles) and D2 (white circles) MSNs. Capsaicin was applied for 30 min (black bar) after at least 10 min baseline recording. Capsaicin inhibited PFC-evoked oEPSCs similarly in both D1 (black) and D2 (white) MSNs. Capsaicin enhanced BLA-D1 MSN synapses but inhibited BLA-D2 MSN synapses. Finally, capsaicin inhibited vHipp-D1 MSN synapses and has no effect at vHipp-D2 MSN synapses. **B)** Individual and averaged oEPSCs amplitude experiments 10 min before (baseline) and 25-30 min after capsaicin; D1 MSNs, black circles: PFC n= 3, p=0.0308, BLA n=4, p=0.0188, vHipp n=5, p=0.0010; D2 MSNs, white circles: PFC n=3, p=0.0135, BLA n=3, p=0.0087, vHipp n=3, p=0.0109 paired t-test). **C)** Summary bar histogram of the maximal effects of Capsaicin on oEPSCs at identified inputs onto D1 (PFC n= 3, BLA n=4, vHipp n=5, F_(input 2,9)_=157.4, p<0.0001, one-way ANOVA) and D2 MSNs (PFC n=3, BLA n=3, vHipp n=3, F_(input 2,6)_=8.586, p=0.0174, one-way ANOVA). **D)** Schematic view of the relative weight and effects of TRPV1R on oEPSCs at PFC, BLA and vHipp inputs onto D1 and D2 MSNs. n represents the number of mice. All values are represented as mean ± SEM, * p<0.05.

The presynaptic localization of CB1R at excitatory inputs in the NAc core has long been established (Robbe et al., 2001) and synaptic release of eCB triggers long-term depression (eCB-LTD, Robbe et al., 2002). In mice, postsynaptic TRPV1R colocalize with presynaptic CB1R in the NAc core (Micale et al., 2009) and postsynaptic TRPV1R can trigger LTD on D2 MSNs (Grueter et al., 2010; Neuhofer et al., 2018). In support of these results, we observed that capsaicin altered the frequency of spontaneous’ EPSCs but did not reduce their amplitudes in D1 MSNs and found opposite effects in D2 MSNs (Figure 7-1).

Taken together, these results reveal that while CB1R are uniformly expressed across input pathways onto D1/D2 MSNs, TRPV1R modulate excitatory inputs to the NAc core in a pathway- and cell-type specific manner.

### Cell-type specific endocannabinoid-mediated LTD at amygdala synapses

In the last series of experiments, we focused on the BLA-NAc core synapses. These synapses are instrumental to reward-seeking behavior (Ambroggi et al., 2008) and have a role in positive emotional valence (Beyeler et al., 2018). Our present observation that both TRPV1R and CB1R modulate synaptic transmission in a cell-type specific manner raised the possibility that the eCB interplay controls the polarity of activity-dependent plasticity at BLA-NAc synapses.

In the striatum, the principal form of eCB-mediated synaptic plasticity is long-term depression (LTD(Robbe et al., 2002)(Gerdeman et al., 2002). We first compared the expression of eCB-LTD at BLA-D1 and -D2 synapses. Using the canonical protocol that elicits eCB-LTD in NAc MSNs (Robbe et al., 2002) we observed a robust LTD in D2 but not D1 MSNs following optical stimulation (Figure 8). This is, to our knowledge, the first evidence that eCB-LTD can be optogenetically-induced at a single set of identified synapses. While this experiment can be interpreted as a lack of eCB-LTD at BLA inputs onto D2 MSNs, the current pharmacological characterization (Figure 6 & 7) and previous work in the NAc core (Neuhofer et al., 2018) led us to test the possibility of another, more complex scenario. Based on the enhancing effects of the TRPV1R agonist at BLA-D1 synapses, we hypothesized that tetanus-induced eCB release (presumably anandamide) and the ensuing potentiation masked/prevented the induction of LTD. In support of this idea we observed that in the presence of a TRPV1R antagonist (capsazepine, 10 μM) the otherwise inefficacious protocol triggered a large LTD (Figure 8C-D). The mirror situation was observed at BLA-D2 synapses. There, LTD was converted to LTP in the presence of the TRPV1R antagonist (Figure 8E-F), a result compatible with the current finding that TRPV1R are excitatory on BLA-D2 synapses (Figure 7). We finally controlled for the role of CB1R in BLA-D2 LTD. Bath-application of the CB1R specific and neutral antagonist NESS (1 μM) reduced post-tetanic depression but did not prevent LTD (Figure 8E-F).

**Figure 8:**
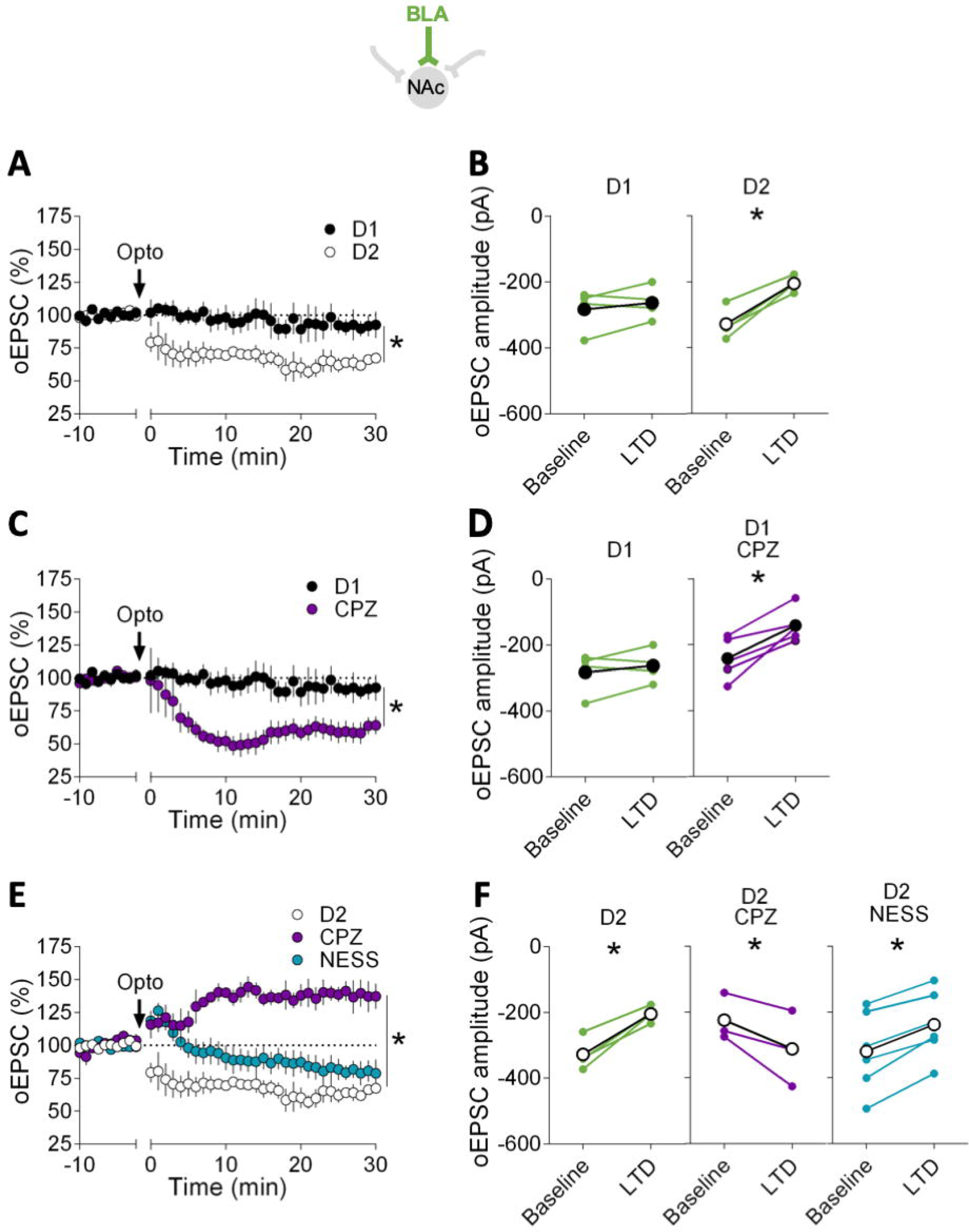
TRPV1R delineates cell type-specific LTD at amygdala synapses in the NAc core. **A)** Optical stimulation of BLA inputs (10 Hz for 10 min) induced a a robust LTD in D2 (white circles, n=4) but not in D1 MSNs (black circles, n=4). **B)** Individual and averaged oEPSC amplitude before (baseline) and 30 min after LTD induction (LTD) in D1 and D2 MSNs (D1 MSNs, p=0.3736, D2 MSNs, p=0.0072, paired t-test). **C)** Antagonism of TRPV1R (capsazepine, CPZ 10 μM) allowed for the induction of LTD at BLA-D1 synapses (purple, n=6; compared to control same as A). **D)** Individual and averaged oEPSCs amplitude before (baseline) and 30 min after LTD induction (LTD) in D1 MSNs with or without CPZ (D1 CPZ, p=0.0056 paired t-test). **E)** Antagonism of TRPV1R converted LTD to LTP at BLA-D2 synapses (purple, n=3). Bath-application of the CB1R specific neutral antagonist NESS 0327 (1 μM) altered without blocking LTD at BLA-D2 synapses (blue, n=7). **F)** Individual and averaged oEPSCs amplitude before (baseline) and 30 min after LTD induction (LTD) in D2 MSNs with or without CPZ or NESS 0327 (D2 CPZ, p=0.0462, NESS, p=0.0039 paired t-test). n represents the number of mice. All values are represented as mean ± SEM, * p<0.05 paired t-test.

## DISCUSSION

Our principal findings are that in the adult mouse NAc, 1/ the hierarchy of excitatory inputs depends on the identity of the postsynaptic target MSN and on circuit specific feedforward inhibition and 2/ the eCB system endows excitatory circuits of the NAc with pathway-specific plasticity.

### Differential intrinsic properties of adult NAc core MSNs

The data show that D1 and D2 MSNs in adult NAc core have distinctive intrinsic properties. Compared to D2, D1 MSNs had a low rheobase, a more depolarized membrane potential and exhibited a higher propensity to trigger action potentials in response to depolarizing current injections. Thus, adult D1 are more excitable than D2 MSNs.

Subtype specific morphological properties (e.g. soma size or dendritic arborization, (Gertler et al., 2008) and differences in membrane biophysical properties such as membrane resistance, capacitance, or the expression of voltage dependent calcium channels, may explain these dissimilarities (Nisenbaum et al., 1994; Hernández-López et al., 1997, 2000).

Divergent intrinsic properties have already been described in in juvenile and adolescent mice, however at these stages D2- are more excitable than D1 MSNs (Kreitzer and Malenka, 2007; Gertler et al., 2008; Ma et al., 2012; Planert et al., 2013; Cao et al., 2018). While many factors (e.g. recording site, sample size, the presence or absence of TTX, the transgenic mouse line used, the size of the soma or the extent of dendritic arborization) may explain this discrepancy, it is highly probable that age is the determining factor. In keeping with this idea, in rat NAc core, prepubertal rats and adult rats display different properties (Belleau and Warren, 2000; Zhang and Warren, 2008; Kasanetz and Manzoni, 2009). Additionally, in both rats and mice, NAc dopamine receptors undergo significant developmental changes during adolescence (Andersen et al., 1997; Andersen and Teicher, 2000).

The subtype specific evolution of excitability in juvenile/adolescent and adults MSNs may be due to differences in membrane resistance and conductance. In support of this possibility, early in development MSNs have very high membrane resistance and do not express inward rectifying potassium channels (Tepper et al., 1998; Belleau and Warren, 2000). The time-scale of this physiological maturation parallels the morphological development of MSNs and dendritic arborization, spine formation and synaptogenesis continue until the end of the first postnatal month (Tepper et al., 1998; Butler et al., 1999).

### The hierarchy of excitatory afferents depends on MSNs’ subtype in adult NAc core

Pathway-specific opto-stimulation of excitatory inputs in the NAc core revealed that there is a hierarchy of synaptic inputs that depends on the cellular identity of the target MSNs. Most notably, the BLA and the PFC are the main source of synaptic excitation on D1 and D2 neurons, respectively. Similarly, BLA and PFC afferents trigger action potentials with a high probability in D1 and D2 MSNS, respectively.

Differences in the number of fibers per afferent or the amount of glutamate release per fibers may underlie this functional hierarchy. Based on the recent study of Li and colleagues (Li et al., 2018) that showed similar PFC, BLA and vHipp innervations of D1 and D2 neurons, we favor the second proposition.

### Pathway and cell type specific feedforward inhibition

Feedforward circuits in the NAc core allow input-evoked activation MSNs to be tweaked by GABAergic monosynaptic innervation from neighboring interneurons. Anatomical studies have shown that MSNs have dense axonal collaterals that remain within the NAc (Chang and Kitai, 1985; Pennartz and Kitai, 1991; Burke et al., 2017). We found that D1 and D2 MSNs receive different levels of feedforward inhibition. Irrespective of cell types, feedforward inhibition is minimal on vHipp and maximal on BLA afferents. In the PFC pathway, feedforward inhibition is specific to cell type: it is minimal on D2 and maximal on D1 MSNs. Thus, it is possible that the recruitment of interneurons is pathway selective or that specific interneurons preferentially inhibit D1 or D2 MSNs. In keeping with this idea, a recent study shows that vHipp afferents preferentially activates PV^+^ interneurons (Scudder et al., 2018).

We also observed that the vHipp-NAc pathway has the unique ability to trigger action potentials in both D1 and D2 MSNs in the presence of “physiological" feedforward inhibition. This may be due to the hyperpolarized resting membrane potential of MSNs associated with an omnipresent inhibitory circuit. In any case, this finding is consistent with the literature: the vHipp pathway is the only one able to trigger action potentials in the NAc shell (Britt et al., 2012) and these very afferents gate the PFC’s output (O’Donnell and Grace, 1995; Goto and O’Donnell, 2001).

### Input and synapse specific function of endocannabinoids

The present data indicate that functional CB1R are present at all synapses tested, albeit with differences in the amount of CB1R-mediated inhibition. The situation was far more complex with TRPV1R. Indeed, TRPV1R agonism inhibited PFC-evoked oEPSCs in both D1 and D2 MSNs but lead to opposite and MSN subtype-specific effects at the BLA pathway: BLA-D1 were enhanced while BLA-D2 synapses were inhibited by capsaicin. TRPV1R specifically inhibited vHipp-D1 synapses and had no effect on vHipp-D2 synapses. Pharmacological activation of TRPV1R suppresses or facilitates neurotransmitter release into the dorsal striatum according to their pre- or postsynaptic location (Musella et al., 2009) and our results with sEPSCs also suggest a differential expression of TRPV1R at BLA-D1 and BLA-D2 synapses. Indeed, capsaicin modified the frequency but not the amplitude of sEPSCs in D1 MSNs and vice-versa in D2 MSNs. These data are compatible with presynaptic TRPV1R at BLA-D1 and postsynaptic TRPV1R at BLA-D2 synapses. Noteworthy, Grueter et al. also reported a postsynaptic localization of TRPV1R at D2 MSNs (Grueter et al., 2010).

At the BLA-NAc pathway, TRPV1R and CB1R modulate synaptic transmission in a cell specific manner: LTD could be induced in D2 but not in D1 MSNs in agreement with a previous report where the identity of MSNs was not ascertained (Grueter et al., 2010). At first glance, the results may be taken as an indication that BLA-D2 MSNs are incapable of expressing eCB-LTD. However, the current pharmacological characterization and previous work (Neuhofer et al., 2018) led us to hypothesize that the tetanus-induced eCB (presumably anandamide), activates TRPV1R and that the ensuing potentiation masked/prevented LTD. In support of this idea, we observed that in the presence of a TRPV1R antagonist, the previously inefficient protocol triggered a large LTD. The mirror situation was observed at BLA-D2 synapses where LTD was converted to LTP in the presence of the TRPV1R antagonist.

In summary, our study reveals the cell-type-specific synaptic organization of hippocampal, amygdala and prefrontal inputs to the PFC. It is tempting to speculate that pathway and cell specific synaptic strengths correlate with distinct functions and associated behaviors. Future work will determine if the PFC-D2 “dominant pathway” is preferentially engaged in executive functions and the role of the BLA-D1 pathway in emotional behaviors. In this context, the observation that two subpopulations of BLA glutamatergic neurons project to D1 or D2 MSNs NAc core to generate emotional responses of opposite valence is an indication of similar parallel dual control circuits at PFC- and/or vHipp-NAc pathways (Shen et al., 2019).

Moreover, the data highlight the versatility of the endocannabinoid system in shaping activity dependent synaptic plasticity at BLA-NAc circuits and cortical-limbic circuits in general.

In conclusion, our experiments reveal a high degree of synapse and circuit specificity in the adult NAc core and illustrate how endocannabinoids contribute to pathway-specific synaptic plasticity.

## Supporting information

Extended data Deroche

## ACKNOWLEDGMENTS

The authors are grateful to Dr. Pascale Chavis and members of the Chavis-Manzoni laboratory for helpful discussions, Dr. Andrew Scheyer for critical reading of the manuscript, Steeve Maldera for experimental help, Dr. Daniela Neuhofer for initiating the project and to the National Institute of Mental Health’s Chemical Synthesis and Drug Supply Program (Rockville, MD, USA).

## AUTHOR CONTRIBUTIONS

MD, OL and OJM designed research; MD and OL performed research; MD and OL analyzed data; MD and OJM wrote the paper.

