## Extended data Deroche for "Cell-type and endocannabinoid specific synapse connectivity in the adult nucleus accumbens core"

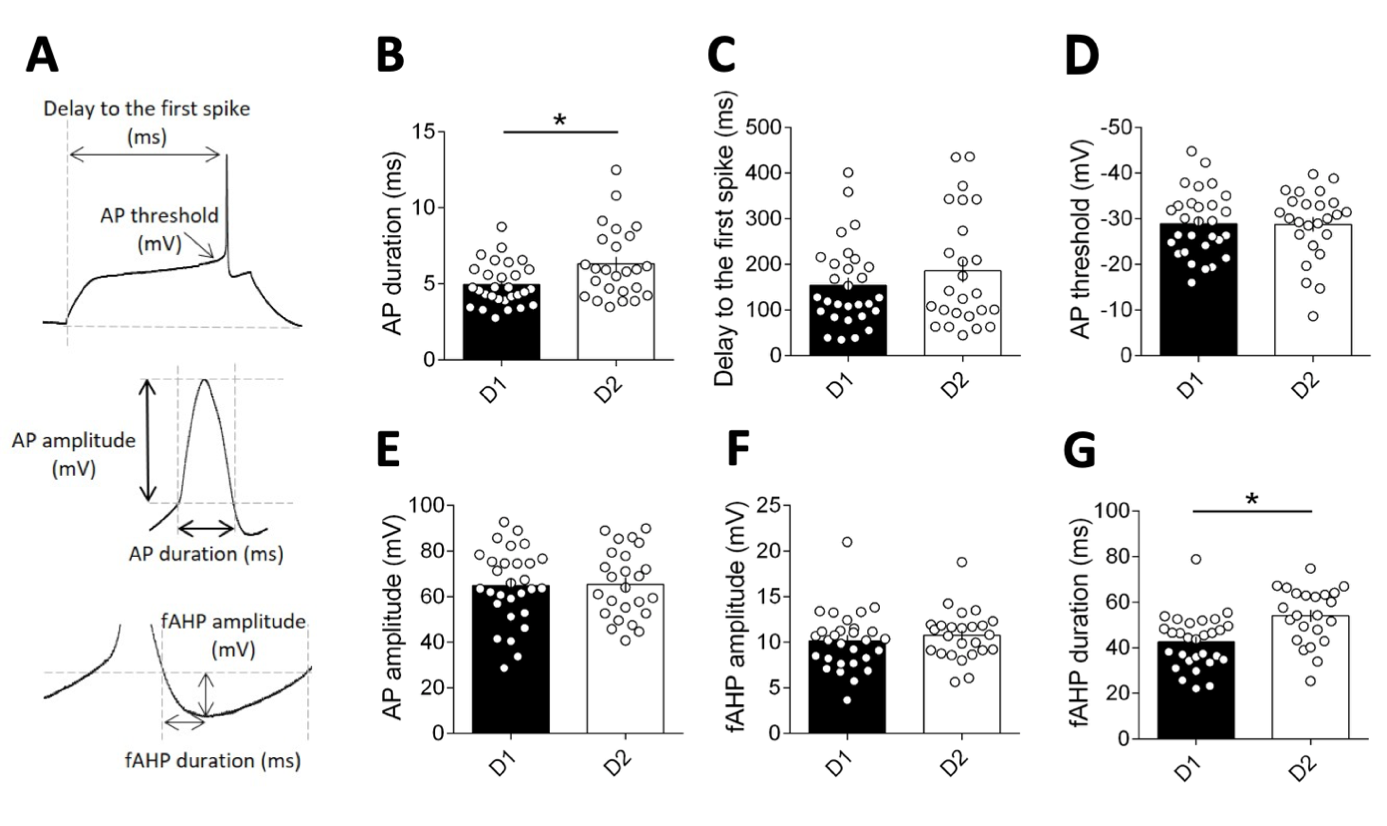
**Figure 1-1: Action potential properties: AP duration and fAHP duration varies by MSN subtype**

**A)** Example of individual AP evoked by depolarizing current injection, indicating delay to the first spike, AP threshold, AP amplitude, AP duration, fAHP amplitude and fAHP duration metrics. **B)** AP duration is shorter in D1 MSNs compared to D2 MSNs (D1 MSNs n=30; D2 MSNs n=26, p=0.0354, Mann-Whitney U test). The following action potential properties did not vary between the MSN subtype: **C)** delay to the first spike. **D)** AP threshold. **E)** AP amplitude. **F)** fAHP amplitude. **G)** fAHP duration is increased in D2 MSNs compared to D1 MSNs (p=0.0009 Mann-Whitney U test). Scatter dot plots represents one cell. All values are represented as mean ± SEM.

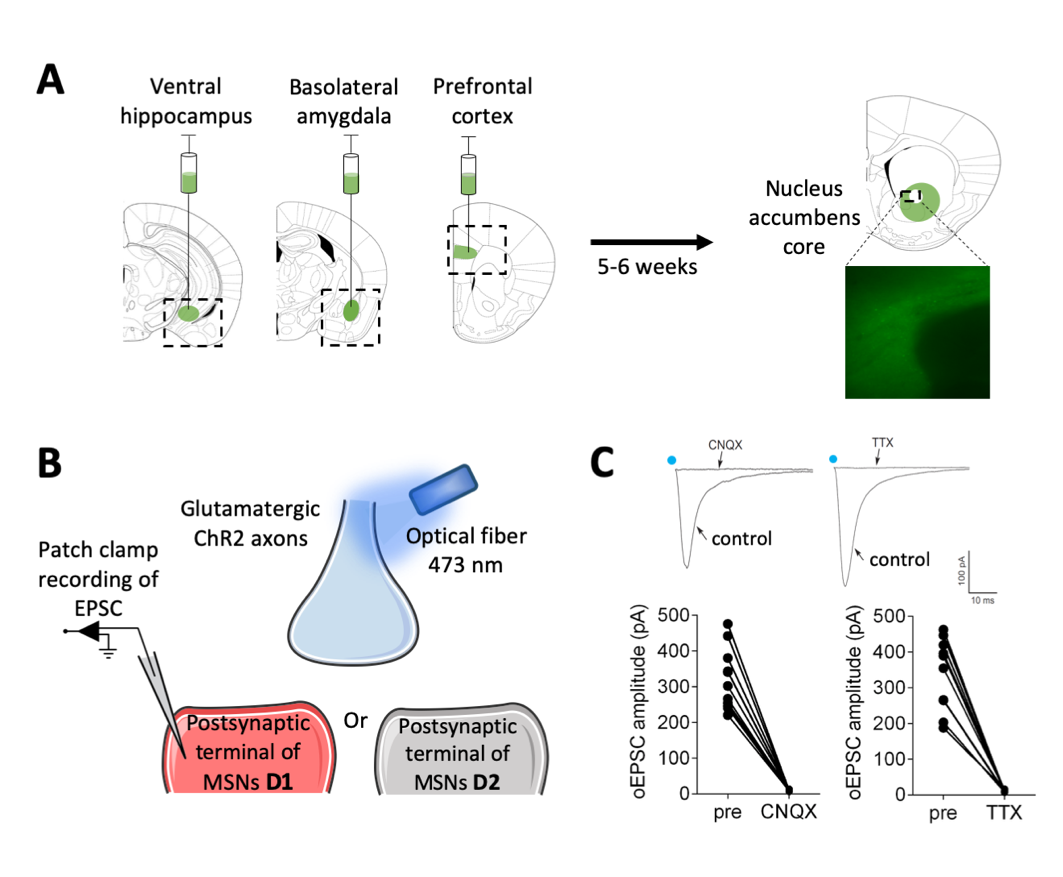

**Figure 2-1: Pathway specific evoked excitatory postsynaptic currents (EPSCs) in the Nac core following optical stimulation of ventral hippocampal, basal amygdala or prefrontal cortex inputs**

**A)** Representative coronal brain slices showing expression of ChR2-eYFP (green) following injections of (0.25 µL) AAV9.CamKIIa.hChR2(H134R)-eYFP.WPRE.hGH (Addgene26969P; 1.98 x 10 GC/mL)) in the ventral hippocampus (vHipp), basolateral amygdala (BLA) or prefrontal cortex (PFC) (left). Image of ChR2-EYFP expressing axons from principal (i.e. CamKII expressing) cortical neurons (right). **B)** Illustration of the experimental set up. Synaptic terminals expressing ChR2-EYFP were stimulated with a 473 nm laser coupled to an optical fiber placed 350 µm from the recording area. Recordings of Optically-evoked EPSCs were recorded in the whole-cell patch-clamp configuration in medium spiny neurons (MSNs) from the NAc core. **C)** Inward currents evoke by light stimulation in NAc core MSNs. CNQX (20 µM) and TTX (1 µM) completely prevented evoked currents following optical stimulation of PFC inputs, showing that the oEPSCs depended on presynapitc and post-synaptic glutamate ionotropic AMPAR receptor-mediated currents. Individual oEPSCs amplitude experiments before (pre) and after CNQX (n=10) or TTX (n=9). Blue dots indicate optical stimulations.

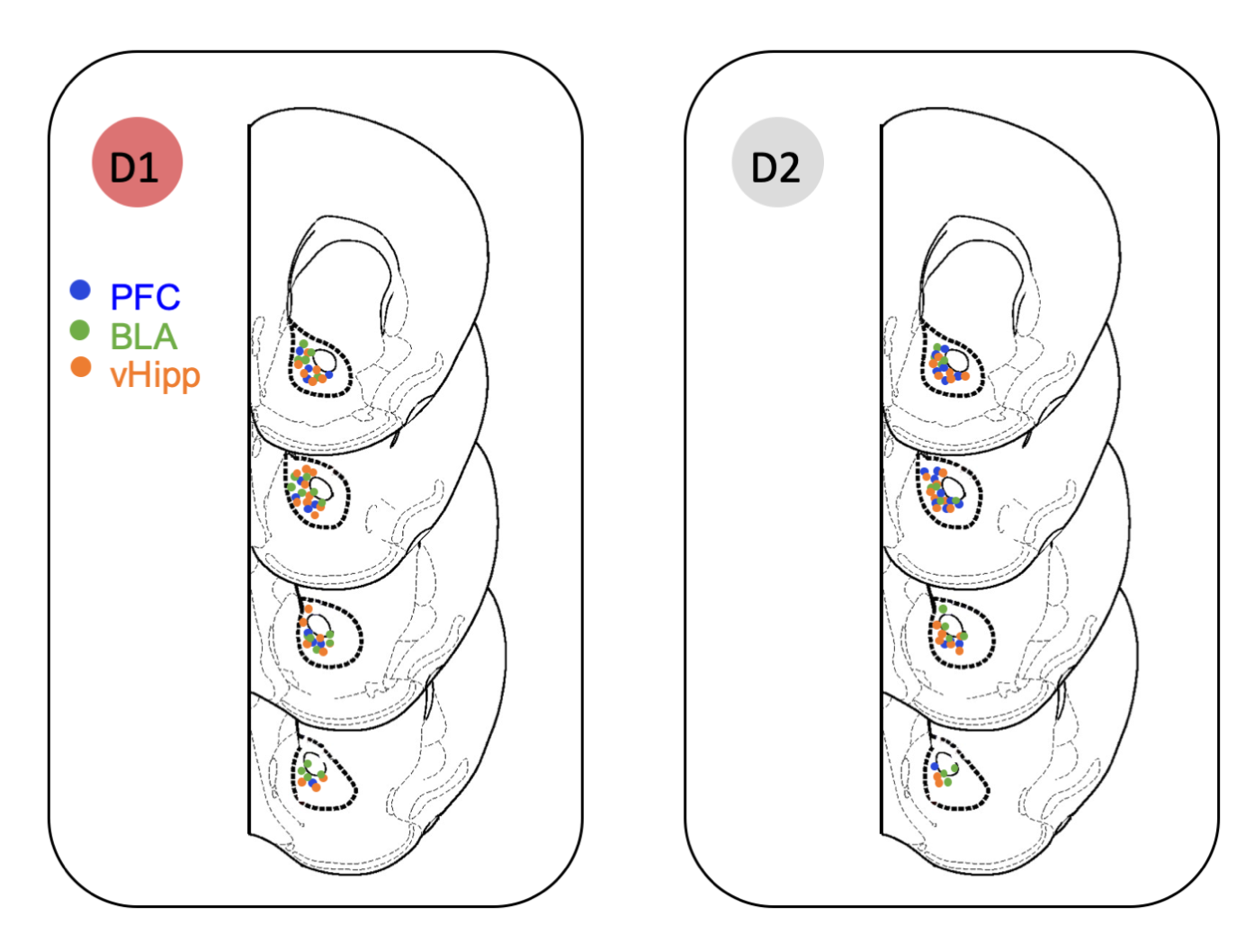
**Figure 2-2: Location of whole-cell patch-clamped MSNs sorted by subtype in nucleus NAc core of Drd1-tdTomato transgenic mice**

“D1” represent recordings from fluorescently-labeled D1-positive MSNs and “D2” from non-fluorescently-labeled MSNs.

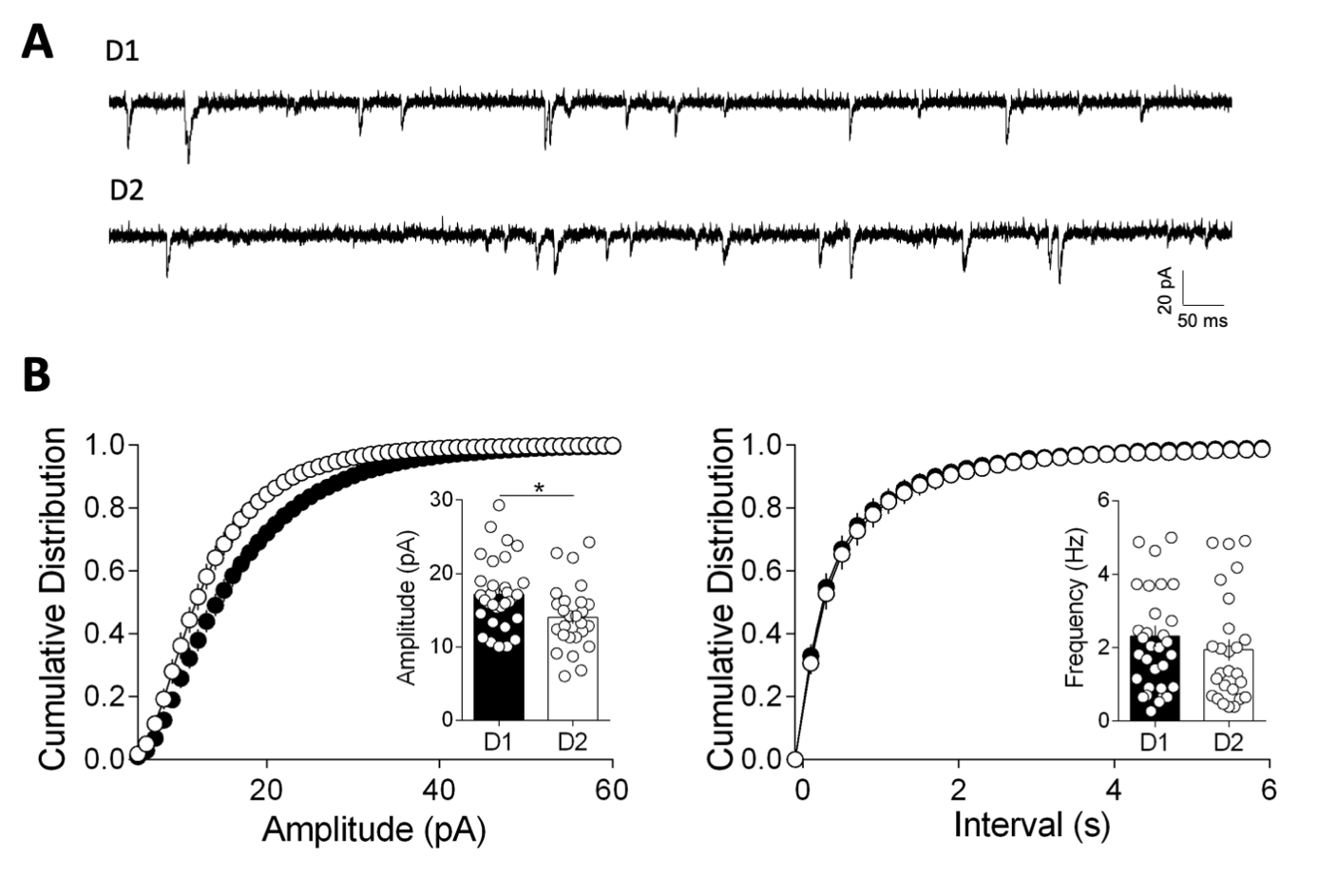

**Figure 5-1: Cell-type-specific differences in amplitude of sEPSCs in MSNs**

**A)** Representative traces of whole-cell voltage clamp recording AMPAR event sEPSC from in D1 and D2 MSNs. **B)** Left, Cumulative probability plot of measured sEPSC event amplitudes. Summary bar chart shows mean AMPAR event sEPSC amplitude (D1 MSNs: n=32; D2 MSNs: n=26, p=0.0146, Mann-Whitney U test). Right, Cumulative probability plot of interval between measured sEPSC events. Scatter dot plots represents one cell. Summary bar chart shows mean AMPAR event sEPSC interval.

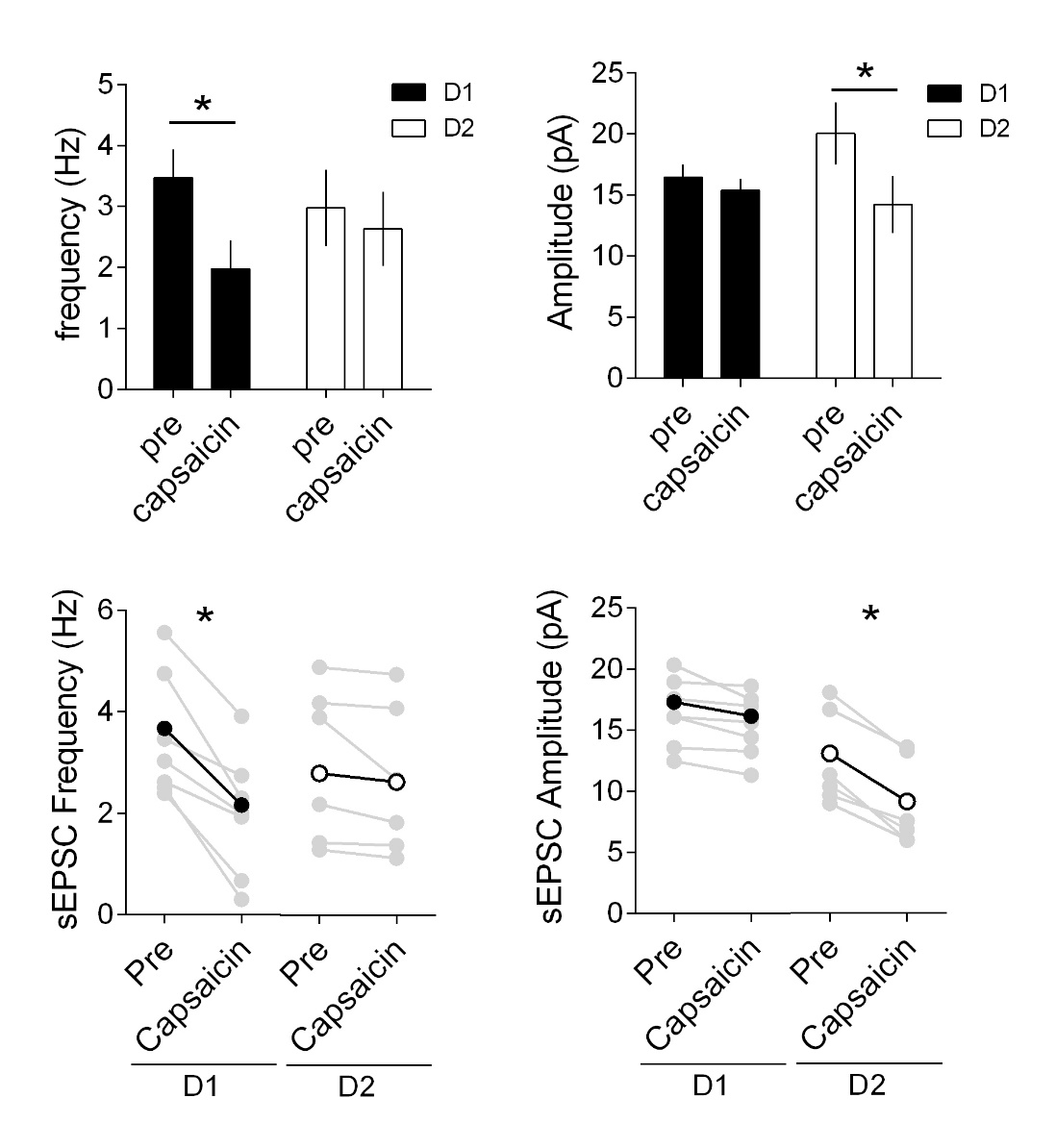

**Figure 7-1: Capsaicin alters the frequency of sEPSC in D1 MSNs and amplitude in D2 MSNs**

Mean sEPSC frequencies (left) and amplitudes (right) in D1 (black, n=7) and D2 (white, n=6) MSNs before and after capsaicin. All values are represented as mean ± SEM, * p<0.05 paired t-test.
